# Sensory Loss Enhances Multisensory Integration Performance

**DOI:** 10.1101/483586

**Authors:** Moa G. Peter, Danja K. Porada, Christina Regenbogen, Mats J. Olsson, Johan N. Lundström

## Abstract

Auditory and visual sensory loss has repeatedly been shown to alter abilities in remaining sensory modalities. It is, however, unclear whether sensory loss also impacts multisensory integration; an ability that is fundamental for the perception of the world around us. We determined effects of olfactory sensory deprivation on multisensory perception by assessing temporal as well as semantic aspects of audio-visual integration in 37 individuals with anosmia (complete olfactory sensory loss) and 37 healthy, matched controls. Participants performed a simultaneity judgement task to determine the temporal binding window, and a multisensory object identification task with individually degraded, dynamic visual, auditory, and audio-visual stimuli. Individuals with anosmia demonstrated an increased ability to detect multisensory temporal asynchronies, represented by a narrowing of the audio-visual temporal binding window. Furthermore, individuals with congenital, but not acquired, anosmia demonstrated indications of greater benefits from bimodal, as compared to unimodal, stimulus presentation when faced with degraded, semantic information. This suggests that the absence of the olfactory sense alters multisensory integration of remaining senses by sharpening the perception of cross-modal temporal violations, independent of sensory loss etiology. In addition, congenital sensory loss may further lead to increased gain from multisensory, compared to unisensory, information. Taken together, multisensory compensatory mechanisms at different levels of perceptual complexity are present in individuals with anosmia.

## INTRODUCTION

Sensory deprivation can alter abilities in the remaining senses, often enhancing specific aspects of performance. These altered abilities have been thoroughly studied in isolated sensory modalities, such as auditory or tactile abilities in blind individuals, and the enhanced abilities are often argued to be a compensation for the lost sensory modality. From this compensatory view, it could further be argued that the necessity to integrate information from *multiple* senses to utilize all available input optimally should be even greater when information from one sensory modality is unreliable or completely unavailable. Yet, few studies have explored whether the ability to integrate information from multiple senses, so-called multisensory integration (MSI), is affected by sensory loss.

The literature provides a rich documentation of blind individuals excelling in auditory tasks; primarily related to spatial processing (Lessard et al. 1998; Röder et al. 1999; Voss et al. 2004; Collignon, Voss, et al. 2009), but also to other tasks, e.g., pitch discrimination (Gougoux et al. 2004; Voss and Zatorre 2012). The compensatory abilities displayed by blind individuals are not limited to the auditory domain but extend to tactile (Goldreich and Kanics 2003; Legge et al. 2008) and there are studies showing enhanced performance also on olfactory tasks (Rombaux et al. 2010; Cuevas et al. 2009). In line with these cross-modal compensatory abilities in the blind, deaf individuals have also been shown to outperform hearing controls in tactile (Levänen and Hamdorf 2001) and visual tasks (Bavelier et al. 2006; Dye et al. 2009). These behavioral enhancement effects are commonly stronger in individuals with congenital or early-onset sensory loss, compared to individuals with a sensory loss acquired later in life (in line with the literture, we refer to acquired as well as congenital sensory depriavation as sensory *loss*, despite the fact that individuals with congenital disabilities have not experienced a loss per se; Voss and Zatorre 2012; Gougoux et al. 2004; Voss 2013; Merabet and Pascual-Leone 2010). Although there is strong evidence for cross-modal enhancement effects, also equal (Cornell Kärnekull et al. 2016; Alary et al. 2009; Sorokowska et al. 2018) or even decreased (Zwiers et al. 2001; Lewald 2002; Bolognini et al. 2012) abilities in remaining senses have been observed. The discrepancy of these results can, at least partially, be explained by very heterogeneous patient groups in terms of, e.g., the duration, cause, and severity of sensory loss combined with small sample sizes (not unusual with <10 individuals; Sorokowska et al. 2018; Merabet and Pascual-Leone 2010). The small samples are a logical consequence of the sparseness of individuals displaying complete blind- or deafness in the general population. Individuals with visual sensory impairment or complete auditory sensory loss constitute approximately 0.7 % and 0.2 % of the general Swedish population (Statistiska centralbyrån 2018), while individuals with olfactory sensory impairment (*hyposmia*) and complete olfactory sensory loss (*anosmia*; often used interchangeably with *functional anosmia*: olfactory loss to such an extent that no useful function remains; Hummel et al. 2017) constitute the largest group of individuals with sensory loss, with a prevalence of 20 % and 5 %, respectively (Landis et al. 2004; Brämerson et al. 2004). Despite the high prevalence of anosmia, the number of studies investigating cross-modal behavioral compensation in this population is strikingly low. As of today, studies have focused nearly exclusively on the processing of the remaining *chemical* senses, the gustatory and the trigeminal sense. In sharp contrast to the compensatory abilities often reported in blind and deaf individuals, anosmia has been linked to a reduction in chemosensory abilities (Landis et al. 2010; Frasnelli et al. 2010; Hummel et al. 2003; Gagnon et al. 2014). These negative consequences of olfactory loss have been attributed to the fact that the three chemical senses are strongly interdependent (Frasnelli et al. 2011; Reichert and Schöpf 2018); they form a holistic flavor perception (Small 2012), and are processed in overlapping cortical networks (Lundström et al. 2011). Whether cross-modal compensatory effects of anosmia on *non-chemical* sensory modalities exist remains to be investigated.

Moreover, in addition to investigating cross-modal compensatory effects of anosmia on non-chemical senses, it still needs to be determined whether compensatory effects of sensory deprivation on unisensory processing also extend to multisensory processing. As of today, few studies have addressed the effects of sensory deprivation on MSI. Reasons for this could be an incompatibility of the field of sensory deprivation and the field of MSI, with the former focusing on blind and deaf individuals, whereas the majority of experimental paradigms investigating MSI depend on the integration of visual and auditory inputs. Therefore, studying individuals with anosmia to investigate the effects of sensory loss on MSI provides two strong advantages compared to studying blind or deaf individuals: it facilitates the use of larger samples sizes and enables the use of established audio-visual integration paradigms not suited for blind or deaf individuals.

In general, the integration of complementary information from different senses leads to behavioral benefits, such as enhanced accuracy, improved detection, and reduced response time (Stevenson, Ghose, et al. 2014; Murray et al. 2016; Stein and Stanford 2008). To better utilize sensory integration of remaining senses would therefore be of great compensatory benefit to individuals suffering from sensory loss. Moreover, there is a strong link between olfactory and multisensory processing, with cross-modal and multisensory effects evident in primary olfactory areas such as the olfactory bulb (Czarnecki et al. 2018), olfactory tubercle (Wesson and Wilson 2010), and piriform cortex (Porada et al. 2018), and it was recently demonstrated that multisensory areas *directly* influence olfactory cortex when processing visuo-olfactory stimuli (Lundström et al. 2018). Given this link between early olfactory areas and multisensory processing, altered MSI performance could be expected as a consequence of olfactory loss. Further support for the notion of altered MSI as a consequences of sensory loss is provided by the functional reorganization of cerebral areas linked to MSI processing in sensory deprived animals and humans (Hyvärinen et al. 1981; Bavelier and Neville 2002). Taken together with the cross-modal enhancement effects that have been demonstrated in other sensory loss populations, it is sensible to assume that an increased, rather than decreased, MSI performance would be manifested in individuals with anosmia.

We hypothesized that the loss of olfactory functions has supra-modal compensatory consequences and results in more efficient information processing of multisensory stimuli. We assessed the ability to integrate multisensory information in individuals with anosmia and matched, healthy controls using two well-established audio-visual experimental paradigms that focus on temporal as well as on semantic aspects of audio-visual integration. One task employed perceptually simple audio-visual stimuli to investigate temporal integration of multisensory stimuli. Specifically, we assessed the audio-visual temporal binding window (TBW), a limited time span between two stimuli during which binding of the two into one percept is highly probable (Vroomen and Keetels 2010; Stein and Stanford 2008; Stevenson et al. 2012). The other task focused on integration of more complex, dynamic stimuli (audio and video clips of common objects) in an object identification task. The task was based on the principle of inverse effectiveness, stating that weaker unimodal stimuli (i.e., stimuli more difficult to perceive) lead to stronger integration effects (Stein and Meredith 1993); the stimuli were therefore degraded (overlaid with noise) and presented either uni- or bimodally to investigate the performance gain for multi- as compared to unimodal object presentations. In combination, these established experimental tasks investigating various MSI abilities enabled us to explore a wide spectrum of potential cross-modal compensatory abilities in the form of audio-visual integration in individuals with anosmia.

## METHOD

### Participants

A total of 74 participants in the age span of 18-59 were included in the study: 37 individuals with isolated, non-traumatic, functional anosmia (25 women; mean age 36.7 ± 11.2 years; sample size limited by patient availability) and 37 healthy controls (25 women; mean age 35.9 ± 12 years), matched in terms of sex, age, and educational level. The reported cause of anosmia was congenital (n = 25), with no recollection of ever experiencing odors, and acquired anosmia, subdivided into upper respiratory tract infections (n = 6), idiopathic (n = 5), and allergic reaction (n = 1). The duration of olfactory loss for participants with acquired anosmia was at least 22 months prior to participation in the study (inclusion criteria duration > 1 year, mean duration 10.7 years, SD = 7.7 years). All participants demonstrated normal/corrected-to-normal visual acuity (exclusion criteria binocular visual scuity < 20/40 according to Snellen’s visual acuity evaluation, Snellen 1862) and auditory (exclusion criteria monaural performance of < 50% correct on a computerized whispered voice test, Pirozzo et al. 2003) functions. The existence of a functional sense of smell (control group) or functional anosmia (patient group), was established with Sniffin’ Sticks 16 odor identification test (Hummel et al. 2007; exclusion criteria < 11 for controls and > 7 for functional anosmia; Control: mean score = 13.6, SD = 1.3, range = 11-16; Anosmia: mean score = 4.4, SD = 1.7, range = 1-7; chance level = 4). Participants did not use psychiatric medications or other medications that could affect their sensory functions, and their intake of alcohol and caffeine was restricted prior to testing. All participants provided written informed consent and all aspects of the study were approved by the local ethical review board.

### Procedure

The study consisted of two audio-visual integration tasks. One task assessed temporal congruency perception between auditory and visual stimuli by means of simultaneity judgements. The other task assessed semantic object identification when presented with dynamic auditory, visual, or audio-visual stimuli. The order of the tasks was counterbalanced with a break in between.

#### Simultaneity judgement task

Temporal congruency perception was assessed by a simultaneity judgement task, in which audio-visual stimulus pairs were presented either temporally synchronized or unsynchronized on a computer screen and through headphones (JBL J55, JBL Inc., Los Angeles, CA). Each trial (Figure 1) began with a white crosshair fixation cross centered on a black background for 1-1.5 s (jittered) before the visual stimulus (white circle with an outer diameter of 15 degrees and an inner diameter of 7.5 degrees of visual angle, centred around the fixation cross) was presented for a duration of 16.7 ms (one monitor refresh cycle). The auditory stimulus (1.8 kHz beep, 87 dB) was presented binaurally for a duration of 16.7 ms, either simultaneously with the visual stimulus, or at a stimulus onset asynchrony (SOA) ranging from −300 ms (auditory leading) to +300 ms (auditory lagging) divided in 15 steps (0, ±25, ±50, ±100, ±150, ±200, ±250, ±300 ms). The visual stimulus presentation was followed by a 0.5 s fixation cross before a question mark appeared, indicating the onset of a 2 s long response period during which the participant was to respond “simultaneous” or “not simultaneous” using the left and right arrow key on the keyboard (key assignment randomized across participants); omission trials were removed prior to analysis (8 % of all data; no significant group difference in number of omission trials, *t*(49.4) = 1.17, *p* = .25, *d* = 0.27). The response period was followed by a 0.6 s black screen leading to the start of the following trial.

**Figure 1.**
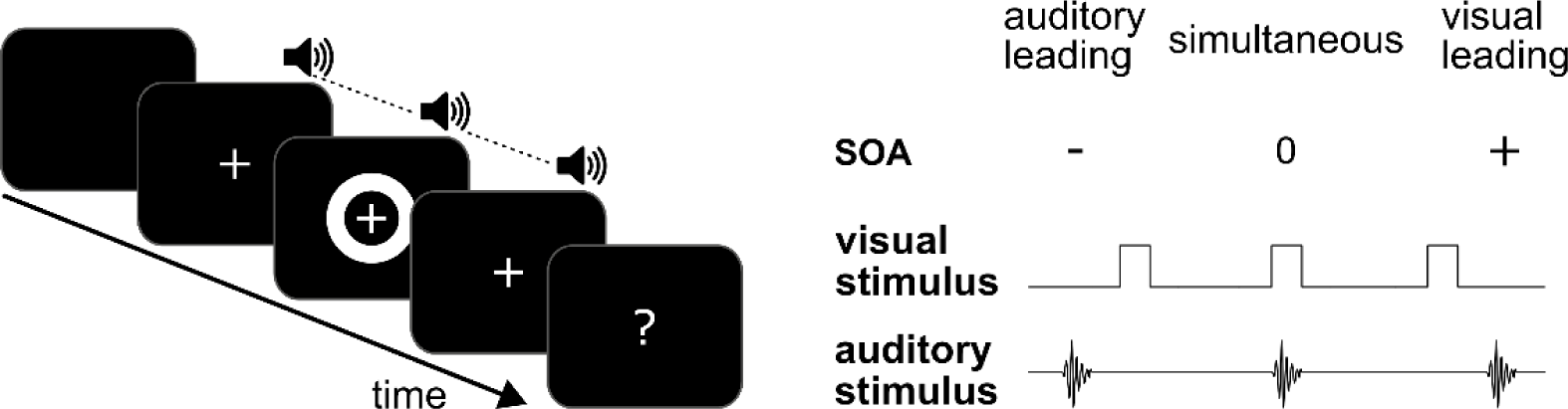
Simultaneity judgement task. The auditory stimulus was presented either before, after, or at the same time as the visual stimulus with a Stimulus Onset Asynchrony (SOA) ranging from −0.3 s to 0.3 s in 15 steps. Participants judged whether the stimuli were presented simultaneously or not.

The task started with five practice trials followed by 200 experimental trials divided into four blocks of equal length with breaks in-between. To minimize response bias, the number of trials in which an asynchrony is assumed perceivable was equalized with the number of trials in which asynchrony is assumed unperceivable, i.e., the experimental trials with SOAs in the range −50 ms to +50 ms were presented with a 2:1 ratio to the experimental trials with SOAs in the range ±100 to ±300 (van Eijk et al. 2008; Vroomen and Keetels 2010; Powers et al. 2009). Stimulus presentation was controlled using PsychToolbox 3.0 for MATLAB 2015b. Auditory temporal accuracy was enhanced using an external soundcard (Scarlett 2i2, Focusrite Inc. High Wycombe, England) and the exact SOAs were measured using a custom built photodiode and auditory signal spectra using PowerLab (ADInstruments, Colorado Springs, CO, USA) to ensure temporal precision in stimulus presentations. Maximum variability of SOA, at any given time, over an average session was ±4 ms.

#### Object identification task

##### Stimuli

Four video clips and four corresponding audio clips clearly representing common, familiar, objects (“wood fire”, “lawn mower”, “popcorn”, and “flopping fish”) with a variety of visual movement and sound types were obtained from Shutterstock (http://www.shutterstock.com). The video and audio clips were edited according to a procedure previously described in detail (Regenbogen et al. 2016; Regenbogen et al. 2018), resulting in stimulus durations of 2000 ms with a 100 ms fade-in and fade-out ramps and a root mean square equalization of −23 dB loudness for audio clips, and a resolution of 720*200 pixels for video clips. In addition to these object depicting clips, “zero information stimuli” of 2000 ms duration, depicting salt and pepper noise (visual; pattern change rate of 30 frames per second) and pink noise (auditory), were created in MATLAB 2015b (The MathWorks Inc., Natic, MA, US).

##### Threshold assessment

Prior to the multisensory object identification task, individual psychophysical thresholds corresponding to a 75 % performance accuracy for each unimodally presented object were assessed by adding noise (visual – salt and pepper; auditory – pink noise; Figure 2) to the stimuli in an adaptive staircase procedure using MATLAB 2015b, as previously described in detail (Regenbogen et al. 2016; Regenbogen et al. 2018). In short, two separate threshold assessment blocks were performed, one for video clips and one for audio clips (counter-balanced order). A fixation cross on a dark grey background was followed by a 2000 ms stimulus presentation (visual stimuli presented centrally on the screen; auditory stimuli binaurally through headphones with a mean loudness level of 67 dB; object order randomized). After the stimulus presentation, an alternative list depicting the four objects and the alternative “nothing” together with assigned keys on the keyboard was presented on the screen for a time period of 1.5 s, during which the participant had to respond. Two consecutive correct responses triggered an increase in noise level and one incorrect response triggered a decrease in noise level. Specifically, to speed up threshold determination, a correct response during the first trial for each object lead to an increase of three noise levels. The noise level increased three levels for each correct response until the first incorrect response (choosing the wrong object, “nothing”, or unanswered trial) reversed the staircase. A total of 15 noise levels existed, ranging from 60 % to 98 % visual noise with 2 % steps, and −7.5 to −25 dB auditory signal-to-noise-ratio (SNR_dB_) with 1.25 SNR_dB_ steps. The procedure stopped after 5 reversals and a mean of the noise levels of the two last reversals was saved as a threshold level. Note that a threshold level was computed for each unimodal object for each individual. Individually degraded stimulus files were created by overlaying the object video and audio clips with noise levels corresponding to the individual’s threshold for that specific object.

**Figure 2.**
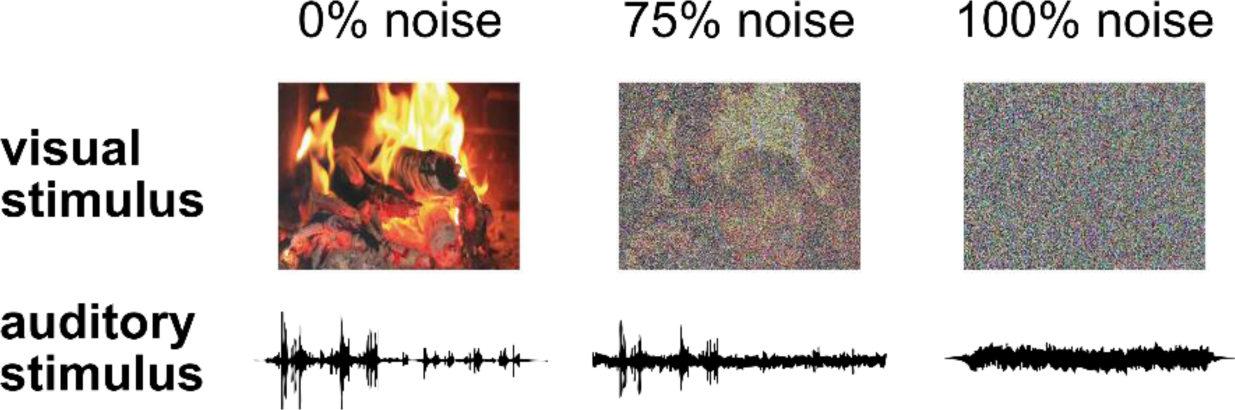
Stimulus examples for the object identification task. Dynamic auditory and visual stimuli of four objects (here represented by the “wood fire” object) were overlaid with noise (visual – salt and pepper; auditory – pink noise) using a staircase procedure to get individually degraded stimuli.

##### Multisensory assessment

In the multisensory object identification task, stimuli consisted of degraded audio and video files representing objects, individually masked based on the threshold assessment to equate the perceived difficulty of unimodal stimuli between all individuals. The degraded audio and video files were presented uni- and bimodally. Specifically, two different approaches to unimodal object presentation were used: one common experimental approach and one more ‘realistic’ approach (Figure 3). In the common experimental approach, only one sense was stimulated at a time with stimuli containing object information, enabling full attention towards that sensory modality: *unimodal experimental setting* (V – visual object, A – auditory object). Although this is the common approach to present unimodal information in MSI-experiments, it is not entirely ecologically valid as we normally receive input to multiple senses at the same time (informative or not). Therefore, in the ‘realistic’ approach, unimodal objects were presented in a bimodal setting, i.e., one sense received object information while the other sense received 100 % noise: *unimodal realistic setting* condition (V_N_ – visual object with auditory noise, A_N_ – auditory object with visual noise). This procedure was adopted to minimize differences in sensory load between the unimodal and the bimodal object presentation (Regenbogen et al. 2016). For the *bimodal object* condition (AV – bimodal audio-visual objects), the video and audio clips were always semantically congruent, i.e., the same object in both sensory modalities.

**Figure 3.**
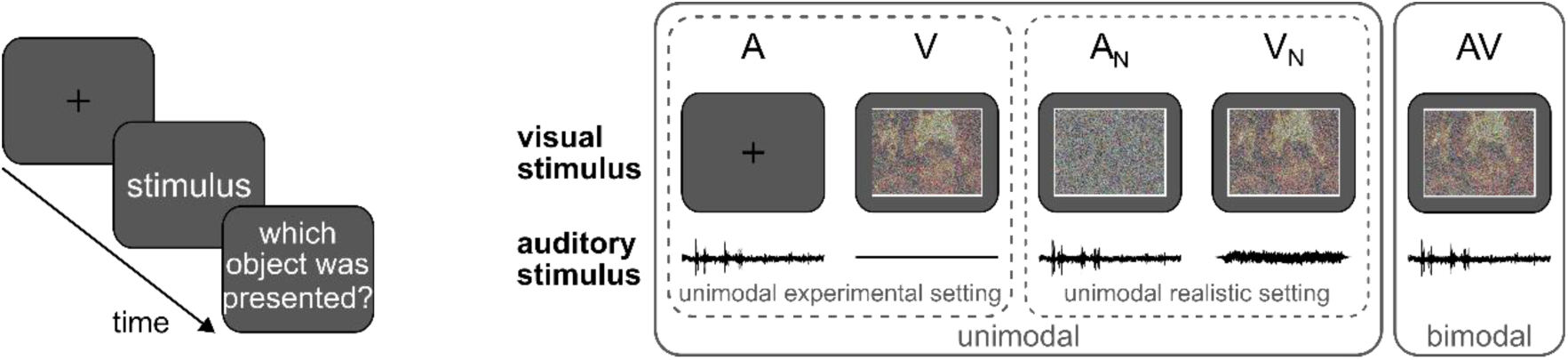
Object identification task: The degraded stimuli, with individual noise levels based on the threshold assessment, were presented either unimodally (A, V; ‘experimental’ setting), as unimodal objects in a bimodal setting (A_N_, V_N_; ‘realistic’ setting), i.e., combined with 100 % noise in the non-informative modality, or as bimodal objects (AV). Participants performed a speeded response task (corresponding to “I have identified the object”) during the 2s stimulus presentation, leading to a 5 choice task (4 objects + “noise” alternative) in which the object was identified.

Each trial began with a fixation cross displayed for a duration of 2 s, followed by stimulus presentation. Participants responded as quickly and accurately as possible by pressing the space bar on the keyboard during the stimulus presentation as soon as they knew which objects was presented (i.e., a maximal response time of 2 s). The key press interrupted the stimulus presentation, giving the participant 2 s to choose one object from the alternative list on the screen (the same five alternatives as during the thresholding, i.e., the four objects and “nothing”). If participants failed to respond before the stimulus presentation was over, no opportunity to choose an object was given and the trial was omitted from subsequent analyses (in total 10.9 % of all data; no significant difference between groups, *t*(71) = 0.03, *p* = .97, *d* < 0.01). Every specific stimulus combination was presented 7 times, resulting in a total of 140 object trials and 35 control trials consisting of pure uni- or bimodal noise. Presentation order was randomized and participants performed a short practice session prior to the experiment, after which they were given the opportunity to ask questions. Stimulus presentation and response collection were controlled by E-Prime 2.0 (Psychology Software Tools Inc., Sharpsburg, PA, USA).

### Data reduction and Analysis

Unless specified, data analyses were performed using the R software package (www.R-project.org) and SPSS (IBM SPSS Statistics for Windows, Version 25.0. Armonk, NY: IBM Corp), and all data exclusion criteria were established prior to analysis based on previous publications (e.g., Regenbogen et al. 2016; Powers et al. 2009)

#### Simultaneity Judgement

Simultaneity perception (the percentage of “simultaneous” responses) was computed for each SOA and participant. Data from two individuals with anosmia and one control subject were removed prior to analysis because they exhibited a mean of less than 75 % simultaneous responses for the trials with SOAs in the span of 0 to ±50 ms, indicating non-compliance with the task. This exclusion criteria was established based on visual inspection of a subset of the data. To assess potential difference between groups in overall attention, response time (RT) over all SOAs was compared between groups with a Welch’s *t*-test, to account for potential differences in variance; effect size estimates are given by Cohen’s *d*.

To assess differences related to perception of simultaneity over groups and SOAs, repeated measures analysis of variance (rmANOVA; within subject factor SOA, between subject factor group) were conducted. Greenhouse-Geisser correction of degrees of freedom was applied if the assumption of sphericity was violated, according to Mauchly’s test; effect size estimates are inferred from partial eta-square (*η*_*p*_^*2*^).

Potential differences between groups in their audio-visual temporal binding window (TBW), i.e., the time span of SOAs during which the auditory and visual stimuli are perceived as simultaneous, were assessed using three analysis steps. First, for each individual, a Gaussian function was fitted to the reported simultaneity perception, with curve parameters *α* (peak amplitude, limited to a maximum of 100), *β* (point of subjective simultaneity, i.e., the SOA for which peak amplitude is reached), and *γ* (standard deviation), according to Equation 1.

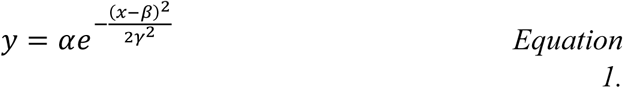

Second, individuals with data for which a Gaussian function could not be fitted were removed from further analysis (two control subjects), leaving a final sample of 35 individuals with anosmia and 34 healthy controls in this specific analyses.

Lastly, the TBW was defined as the span of SOAs during which simultaneity response was more than 75 % of the peak amplitude (*α*), in accordance with previous literature (e.g., Stevenson, Siemann, et al. 2014; Hillock-Dunn and Wallace 2012; Hillock et al. 2011; Powers et al. 2009). Note that by basing the TBW on 75 % of individual peak amplitude instead of 75 % of the theoretical maximal amplitude, analyses were adjusted for potential inter-individual differences in peak simultaneity perception. The TBW was computed for each individual and differences between groups were assessed with a Welch’s *t*-test. Furthermore, to determine potential differences within the anosmia group between individuals with congenital and acquired anosmia, as well as differences between these subgroups and the control group, Welch’s *t*-tests were used. Effect size estimates are given by Cohen’s *d*. Data analyses were performed using the R software package (www.R-project.org) and SPSS (IBM SPSS Statistics for Windows, Version 25.0. Armonk, NY: IBM Corp).

#### Object Identification

##### Threshold assessment

Average unimodal thresholds were computed across objects for each participant. The resulting thresholds (auditory and visual) were compared between groups (Welch’s *t*-test, Bonferroni corrected). To ensure recognition of stimuli during the multisensory assessment, objects that were not identified at a unimodal noise threshold of level four (corresponding to 76% visual noise and −11.25 auditory SNR_dB_) were excluded from subsequent analysis for that participant (11.8% of data).

##### Multisensory assessment

Participants performing with an accuracy below chance level (< .2) for at least one of the five conditions (A, V, A_N_, V_N_, AV) were removed from analysis (five individuals with anosmia, two healthy controls), leaving a final sample of 32 individuals with anosmia and 35 healthy controls for this analysis. Individual trials exceeding mean RT with more than three standard deviations were considered as outliers and excluded from the analysis (five trials, 0.05% of all data). To assess differences in attention between groups, mean RT over all conditions were compared between groups with a Welch’s *t*-test.

Single trial accuracy and RT data was analyzed using a drift diffusion model (DDM; Ratcliff 1978; Voss et al. 2013), a model of cognitive decision making which has been proven successful in modelling decision making in speeded response tasks in both binary and multiple choice tasks (Voss et al. 2013). Recent studies have also demonstrated its usefulness in MSI experiments (Regenbogen et al. 2016; Regenbogen et al. 2018). The inherent trade-off between accuracy and RT measures is accounted for with the use of a DDM, which combines the information from these measures into the model parameters. The DDM represents the two given alternatives in a binary decision process (or the accurate and inaccurate decision when modelling accuracy data) as an upper and a lower decision boundary and assumes that noisy information is continuously accumulated until a decision boundary is reached and a decision is made accordingly. The advantage of the model is its ability to map different cognitive processes related to decision making onto specific variables: the *threshold separation* (*a*; wider separation indicates that more information is needed to reach a decision boundary, i.e., a more conservative decision style), the *non-decision time* (*t*; non-decisional processes such as response execution), the *starting point* or *bias* (*z*; bias towards one option, not applicable when modelling accuracy data, Voss et al. 2013), and the *drift-rate* (*v*; speed at which the decision boundary is reached, a measure of performance/task difficulty), as illustrated in Figure 4.

**Figure 4.**
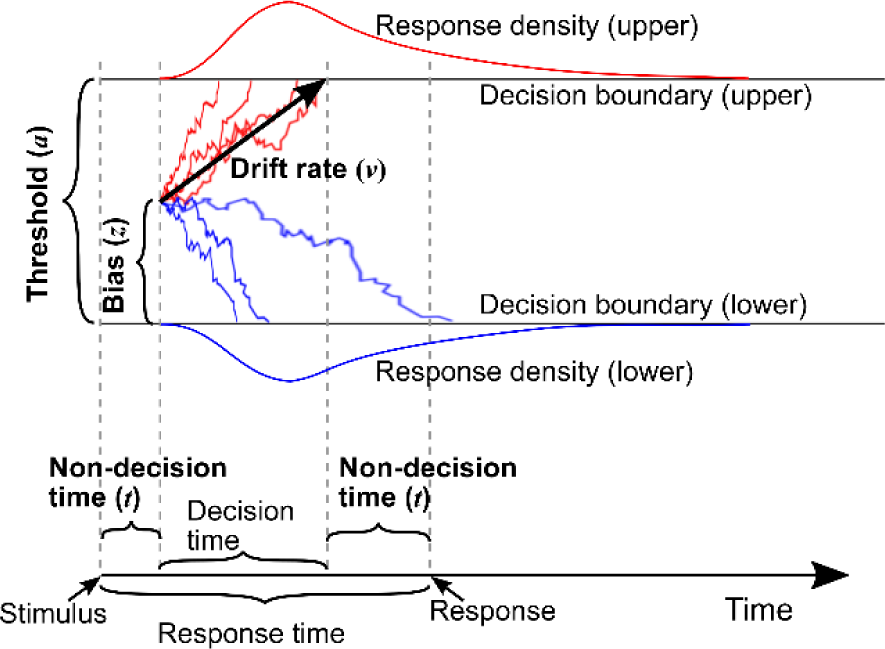
Overview of the drift diffusion model. Evidence is accumulated over time (x-axis) with an average drift-rate **v** until one of two decision boundaries (separated by threshold **a**) is crossed and a response is initiated. The starting point of the decision making process corresponds to the bias **z**. Upper (red) and lower (blue) panels contain density plots over boundary-crossing-times. The flat line in the beginning of the drift-processes, together with the response execution time after reaching a decision boundary, denotes the non-decision time **t** where no accumulation happens.

Here, we used a hierarchical drift diffusion model (HDDM; Wiecki et al. 2013) implemented within an IPython interpreter shell (Perez and Granger 2007); previously proven successful in experimental setups near identical to our current one (Regenbogen et al. 2016; Regenbogen et al. 2018). HDDM uses a hierarchical Bayesian approach in which the individual estimates of model parameters are constrained by group distributions. Based on a DDM likelihood function (Navarro and Fuss 2009), 10 000 posterior samples were drawn using a Markov-chain Monte Carlo algorithm, discarding the first 1000 samples to ensure that model stabilization had been reached. To confirm convergence, multiple runs of the model were performed and the within- and between-chain variance were compared using the Gelman-Rubin *R* statistic (Gelman and Rubin 1992). Furthermore, traces and autocorrelations were visually inspected.

Potential group differences in decision style and response execution were estimated by comparing posterior estimates of the threshold separation (*a*) and non-decision time (*t*) between groups using Welch’s *t*-tests. To investigate potential performance differences between groups and conditions, posteriors of drift-rates (*v*) in the five different conditions (A, V, A_N_, V_N_, AV) were subjected to a rmANOVA (within subject factor Condition, between subject factor Group). Greenhouse-Geisser correction of degrees of freedom was applied if necessary, according to Mauchly’s test of sphericity. Bonferroni corrected *t*-tests between groups for each of the five conditions were subsequently performed.

As a first aim, we assessed whether the experimental manipulation worked, i.e., whether any multisensory enhancement effects (enhanced performance for bimodal as compared to unimodal objects) were displayed. For each individual, the multisensory enhancement of drift-rate, i.e., the performance improvement when presented with bimodal objects as compared to the best performance when presented with unimodal objects, was computed separately for the ‘experimental’ and for the ‘realistic’ setting, according to *ν*_*AV*_ − max(*ν*_*A*_, *ν*_*V*_) and *ν*_*AV*_ − max(*ν*_*AN*_, *ν*_*VN*_), respectively. The enhancements were subjected to one-sample *t*-tests against a no-change value of 0. Thereafter, if significant multisensory enhancements were demonstrated, indicating a functioning experimental manipulation, group differences in enhancements were tested with Welch’s *t*-tests. Similarly, to assess potential differences in multisensory enhancement between individuals with congenital and acquired anosmia, comparison of enhancements between the two subgroups within the anosmia group, as well as differences between the subgroups and the control group, were assessed using independent sample *t*-tests. Effect sizes were given by Cohen’s *d* (*t*-tests) and partial eta-square (*η*_*p*_^*2*^; rmANOVA).

## RESULTS

### Effect of anosmia on audio-visual temporal binding

To assess potential differences in the ability to detect temporal asynchronies of audio-visual stimuli between individuals with anosmia and healthy controls, the percentage of “simultaneous” responses were first computed for each individual and SOA (Figure 5A). A rmANOVA (between subject factor Group, within subject factor SOA) was performed, demonstrating a significant main effect of Group, *F*(1,67) = 9.49, *p* = .003, η_p_^2^ = .12, indicating a difference between individuals with anosmia and controls in perceptual acuity of temporal asynchronies. Furthermore, a significant main effect of SOA was apparent, *F*(5.19, 347.63) = 212.67, *p* < .001, η_p_^2^ = .76, which showed that SOA indeed affects the perceived simultaneity. However, no significant interaction effect between Group and SOA was detected, *F*(5.19, 347.63) = 1.43, *p* = .21, η_p_^2^ = .02.

**Figure 5.**
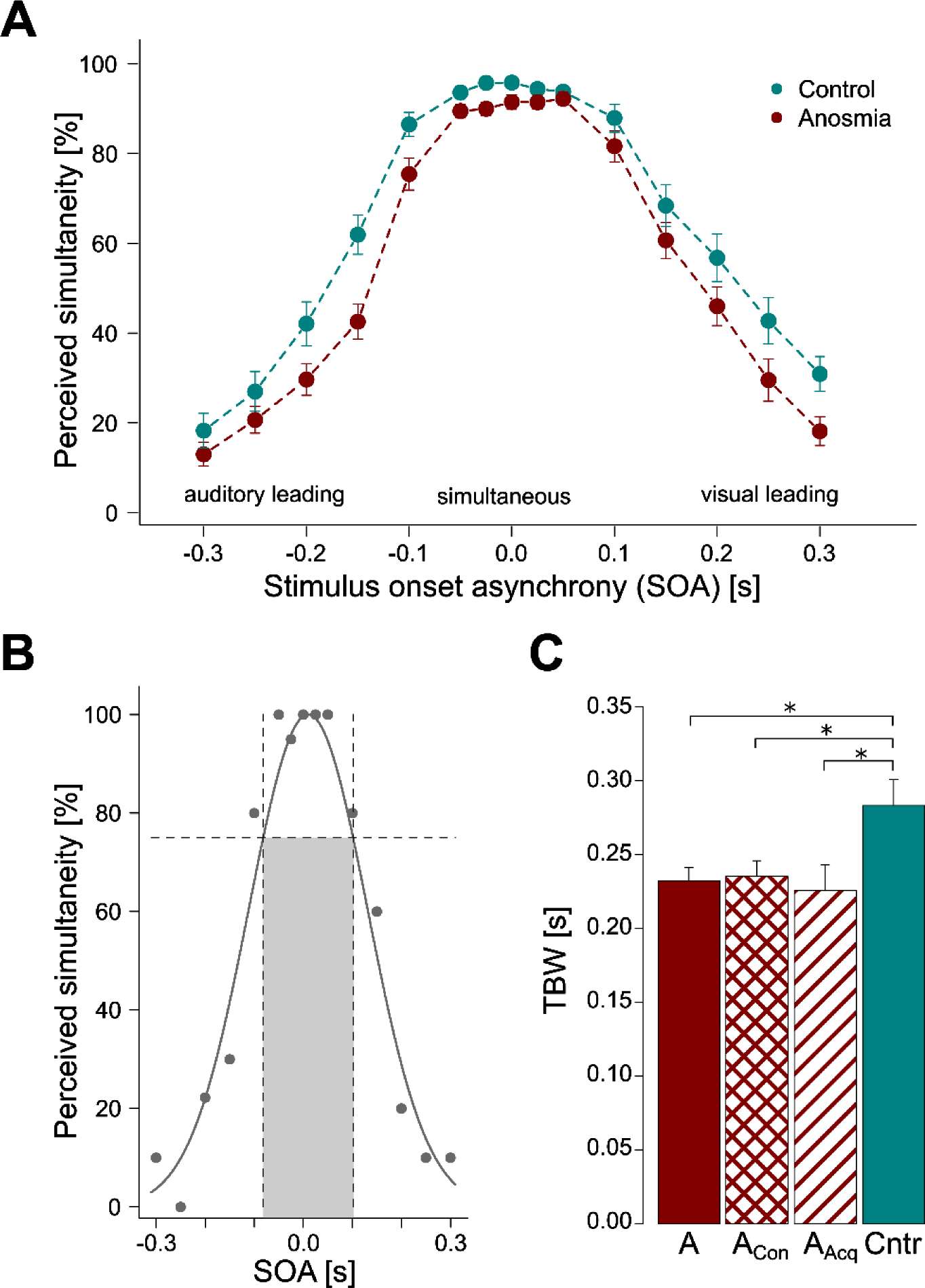
**A)** Group mean proportion of perceived simultaneity for each stimulus onset asynchrony (SOA) level (in seconds). **B)** Example of TBW extraction for a single subject: solid line depicts the fitted Gaussian function, the horizontal dashed line depicts 75 % of maximum simultaneity perception and the vertical lines mark the time span for which the simultaneity perception reach ≥75 % of the maximum simultaneity perception, i.e., the temporal binding window (indicated by shaded area). **C)** Group mean TBW. Individuals with anosmia (A) demonstrated a significantly narrower TBW as compared to controls (Cntr); The narrowing of the TBW was demonstrated by both anosmia subgroups: congenital anosmia (A_Con_) and acquired anosmia (A_Acq_). * = p < .05; error bars indicate standard error of the mean (SEM).

We further determined potential group differences in temporal binding of audio-visual stimuli by a group comparison of the TBW, a threshold value for the temporal integration of stimuli, i.e., an indication of bimodal simultaneity sensitivity. A Gaussian function was fitted to each individual’s response data and the TBW (width at 75 % of maximum) was derived (Figure 5B). The TBW for individuals with anosmia (mean = 232 ms, SD = 54 ms) was significantly narrower than that of healthy controls (mean = 283 ms, SD = 103 ms); *t*(49.4) = 2.57, *p* = .013, *d* = 0.62, i.e., individuals with anosmia demonstrated a better ability to detect small temporal asynchronies in the presentation of audio-visual stimulus pairs (Figure 5C).

Furthermore, potential differences between individuals with congenital and acquired anosmia in temporal binding of audio-visual stimuli were investigated by comparing their obtained TBW. No significant differences were demonstrated between the two subgroups, *t*(17.7) = 0.47, *p* = .64, *d* = 0.17; congenital anosmia: mean = 235 ms, SD = 52 ms; acquired anosmia: mean = 226 ms, SD = 58 ms. Both anosmia subgroups demonstrated a narrower TBW than controls, *t*(51.7) = 2.33, *p* = .024, *d* = 0.59 and *t*(30.8) = 2.31, *p* = .028, *d* = 0.69 for congenital and acquired anosmia, respectively (Figure 5C).

Finally, no significant difference between groups in task attention, operationalized by RT, was found, although there was a statistical trend for faster responses in the control group, *t*(65.4) = 1.71, *p* = .092, *d* = 0.41; Anosmia: mean = 393 ms, SD = 117 ms; Control: mean = 349 ms, SD = 97 ms.

### Effects of anosmia on integration of degraded audio-visual information

#### Threshold assessment

To ensure difficult, but recognizable, stimuli for the multisensory assessment, unimodal thresholds were initially established for each individual and stimuli. There was no significant difference in masking threshold levels for the visual stimuli between groups, *t*(71.3) = 0.19, *p* = .846, *d* = 0.05; Anosmia: mean = 93.24 %, SD = 1.51 %; Control: mean = 93.17 %, SD = 1.67 %; however, the auditory threshold for individuals with anosmia corresponded to a significantly higher SNR as compared to healthy controls, *t*(70.8) = 2.66, *p* = .01 *d* = 0.62; Anosmia: mean = −18.78 SNR_dB_, SD= 2.09 SNR_dB_; Control: mean = −20.16 SNR_dB_, SD = 2.38 SNR_dB_. In the subsequent multisensory assessment, these differences are, however, corrected for by the use of individually determined stimulus masking.

#### Multisensory assessment

Initially, the convergence of the HDDM was examined by visually inspecting the trace (showing no drifts or large jumps), autocorrelation (quickly dropping to zero), and Gelman-Rubin statistic (< 1.02); all values indicated convergence of the model (Wiecki et al. 2013).

Potential differences in overall performance in identifying objects, as measured by drift-rate, between Groups and Conditions (A, V, A_N_, V_N_, AV), were assessed with a rmANOVA. This analysis demonstrated that there was a significant main effect of Condition, *F*(1.66, 107.7) = 57.97, *p* < .001, *η*_*p*_^*2*^ = .47, and a significant interaction between Condition and Group, *F*(1.66, 107.7) = 3.81, *p* < .033, *η*_*p*_^*2*^ = .055; however, there was no main effect of Group, *F*(1,65) = 0.33, *p* = .57, *η*_*p*_^*2*^ = .005. To assess potential group differences for each condition, separate independent samples *t*-tests between groups were computed, revealing no group differences surviving Bonferroni correction for any of the conditions (Table 1).

**Table 1.** Independent sample t-tests comparing drift-rates between individuals with anosmia and controls

| Condition | Drift-rate (v) | | $t$ | $p$ |
| --- | --- | --- | --- | --- |
|  | Anosmia | Control |  |  |
| A | 0.628 (0.5) | 0.36 (0.45) | 2.31 | .024<sup>a</sup> |
| V | 0.645 (0.632) | 0.852 (0.65) | 1.32 | .192 |
| A<sub>N</sub> | 0.512 (0.577) | 0.295 (0.382) | 1.82 | .074 |
| V<sub>N</sub> | 0.678 (0.646) | 0.805 (0.685) | 0.78 | .437 |
| AV | 1.537 (0.557) | 1.394 (0.583) | 0.95 | .345 |

A= Auditory, V= Visual, B=Bimodal, N= Noise presented in the other (V or A) channel. Values within parenthesis indicate SD. <sup>a</sup> denotes that value does not survive Bonferroni correction of 5 comparisons.

We then assessed whether statistical indications of multisensory integration were demonstrated; in other words, whether performance was enhanced for bimodal as compared to unimodal objects, indicated by a significantly larger drift-rate during the bimodal object condition. A one-sample *t*-test indicated significant multisensory enhancement in the ‘experimental’ setting, comparing bimodal objects to unimodal objects; *t*(66) = 12.08, *p* <.001, *d* = 1.48; mean enhancement = 0.48, SD = 0.33. Enhancement effects were also demonstrated in the ‘realistic’ setting, comparing bimodal objects to unimodal objects in ab bimodal setting; *t*(66) = 13.48, *p* < .001, *d* = 1.65; mean enhancement = 0.53, SD = 0.32. This demonstrated a successful experimental manipulation, with multisensory enhancement occurring in both settings.

Having demonstrated successful MSI manipulation, we subsequently assessed whether there was a significant difference between the two groups in multisensory enhancement using Welch’s t-tests. No significant differences between groups in multisensory enhancement were demonstrated in the ‘experimental’ setting, t(64.2) = 1.68, p = .097, d = 0.41 (Figure 6A), nor in the ‘realistic’ setting, t(64.9) = 1.67, p = .099, d = 0.41 (Figure 6B).

**Figure 6.**
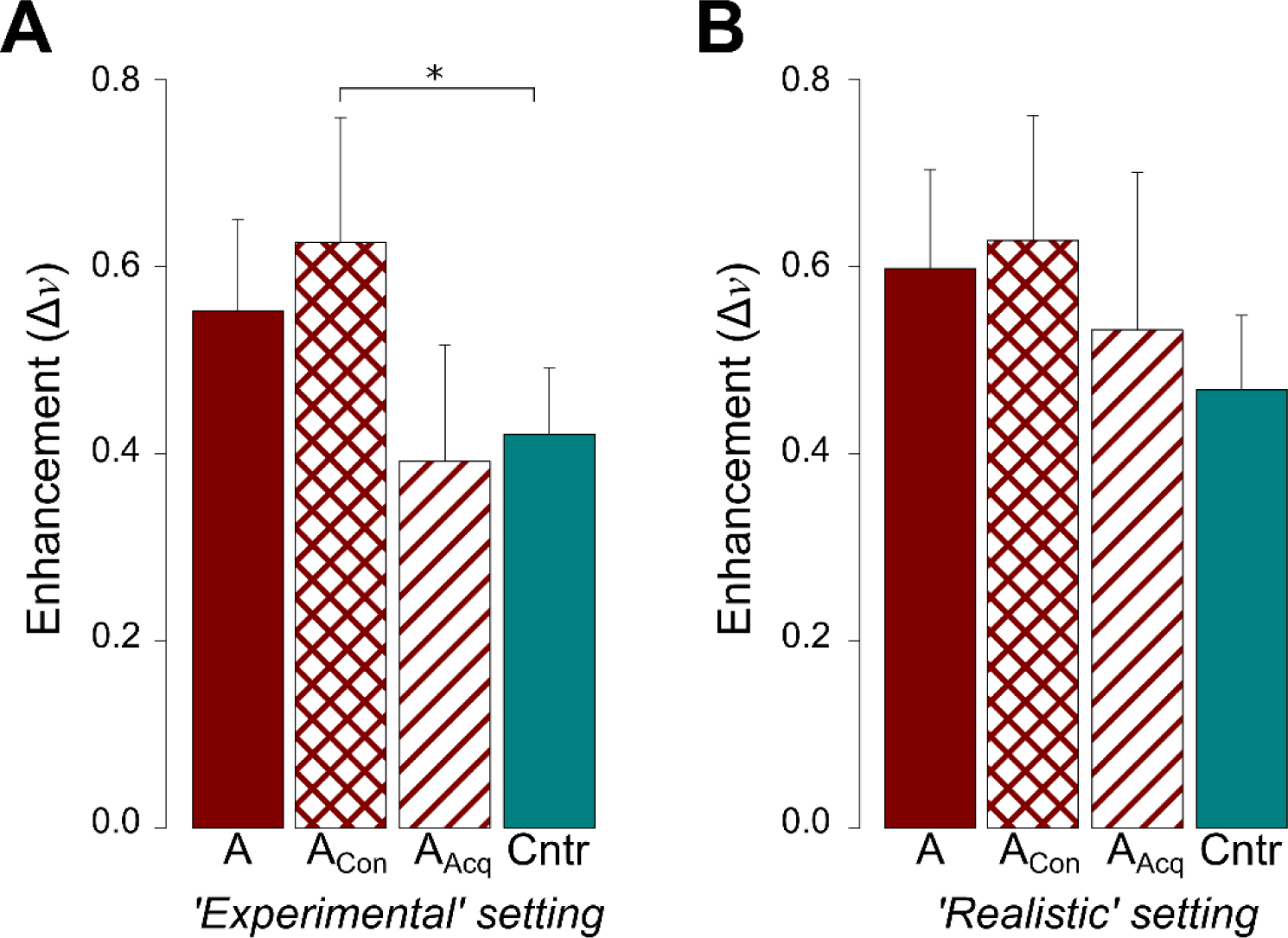
Significant multisensory enhancements in drift rate, derived from the HDDM, were demonstrated, both in the ‘experimental’ setting (ν_AV_ – max(ν_A_, ν_V_)) and in the ‘realistic’ setting (ν_AV_ – max(ν_AN_, ν_VN_)). **A**) ‘Experimental’ setting: No significant difference in multisensory enhancement between individuals with anosmia (A) and healthy controls (Cntr). By dividing the anosmia group into congenital (A_Con_) and acquired (A_Acq_) anosmia, a significantly greater multisensory enhancement was revealed for individuals with congenital anosmia as compared to controls. **B**) ‘Realistic’ setting: No group comparisons revealed significant differences in multisensory enhancement. (* = p < .05 corrected); error bars denote standard error of the mean (SEM).

To investigate potentially different MSI abilities in individuals with congenital and acquired anosmia, the anosmia group was split into respective subgroup. Individuals with congenital anosmia displayed a greater multisensory enhancement, compared to healthy controls, in the ‘experimental’ setting, *t*(54.1) = 2.56, *p* = .013, *d* = 0.67 (Figure 6A). In contrast, no such elevation in multisensory enhancement was demonstrated for individuals with acquired anosmia compared to controls, *t*(16.2) = 0.25, *p* = .806, *d* = 0.09. The comparison between the two anosmia subgroups did not reveal statistically significant differences in performance, even if a statistical trend was demonstrated, *t*(14.5) = 2.08, *p* = .056, *d* = 0.83. In the ‘realistic’ setting, with noise in the non-informative channel, no group differences in multisensory enhancement were demonstrated between any of the three groups, although a statistical trend for greater multisensory enhancement in the congenital anosmia group, as compared to the control group, was demonstrated, *t*(48) = 1.85, *p* = .07, *d* = 0.5 (Figure 6B). Individuals with acquired anosmia demonstrated no indication no statistical differences in multisensory enhancement compared to controls, *t*(16.5) = 0.59, *p* = .563, *d* = 0.2, nor to individuals with congenital anosmia, *t*(18.2) = 0.85, *p* = .407, *d* = 0.32.

Lastly, we assessed whether individuals with anosmia and healthy controls differed in their response behavior and task attention. First, we determined differences related to decision style (threshold separation, *a*) and response execution (non-decision time, *t*). The posterior estimates revealed no statistically significant group difference for threshold separation, *t*(68.8) = 0.24, *p* = .81, *d* = 0.06, nor differences in non-decision time, *t*(64.8) = 0.33, *p* = .74, *d* = 0.08. Finally, potential group differences in attention, operationalized by RT, were assessed by comparing mean RT over all conditions between individuals with anosmia and controls. No statistical differences were demonstrated, *t*(64.4) = 0.035, *p* = .97, *d* = 0.01; Anosmia: mean = 1.3 s, SD = 0.12 s; Control: mean = 1.3 s, SD = 0.15s.

## DISCUSSION

We aimed to determine whether olfactory sensory loss leads to increased multisensory integration (MSI) abilities using two audio-visual binding tasks. Compared to healthy, matched controls, individuals with anosmia demonstrated an enhanced ability to accurately detect short, temporal asynchronies between auditory and visual stimuli of a simple perceptual character. Moreover, individuals with congenital, but not acquired, anosmia demonstrated greater multisensory enhancement than healthy controls in an object identification task with degraded, dynamic stimuli. The difference in multisensory enhancement between individuals with congenital anosmia and controls was, however, only demonstrated when no cross-modal distraction was present for unimodal object identification. The multisensory advantages demonstrated here by individuals with anosmia suggest that olfactory sensory loss leads to cross-modal compensatory abilities in form of enhanced MSI abilities with behavioral advantages more pronounced in individuals with congenital sensory loss.

Individuals with anosmia demonstrated an enhanced ability to detect multisensory temporal asynchronies, which supports the notion that sensory loss affects multisensory perception. Temporal congruency is a key factor influencing multisensory integration, with binding of stimuli only occurring over a limited span of asynchronies (Vroomen and Keetels 2010; Stein and Stanford 2008; Meredith et al. 1987). Deviations in the temporal binding of multisensory stimuli have far-stretching implications. A widened TBW, for example, leads to over-integration of stimuli and has been associated with a variety of clinical conditions, such as autism spectrum disorders, schizophrenia, dyslexia, and obesity (Wallace and Stevenson 2014; Scarpina et al. 2016; Stevenson, Siemann, et al. 2014). In contrast, a narrow TBW has been linked to enhanced performance in both verbal and non-verbal problem solving tasks (Zmigrod and Zmigrod 2016) and individuals with a narrow TBW are less prone to be deceived by multisensory illusions, both when presented with simple perceptual and more dynamic stimuli (Stevenson et al. 2012). These results indicate that a narrow TBW leads to more accurate MSI outcomes and can arguably be viewed as a compensatory mechanism.

Even though the sensory loss literature commonly links congenital or early-onset sensory loss to stronger cross-modal alterations than late-onset sensory loss (Voss et al. 2004), there was no statistical difference in TBW between individuals with congenital and acquired anosmia. The more impactful consequences of sensory loss typically demonstrated by individuals with congenital and early-onset sensory loss are argued to, at least partly, depend on a sensitive period early in life during which sensory experiences shape the development of sensory processing: a period during which a sensory deprivation could increase the potential of cross-modal changes, such as auditory processing in visual cortex (Singh et al. 2018; Voss 2013). The existence of such a sensitive period for multisensory integration in humans has been suggested by Putzar and colleagues (Putzar et al. 2007), who show that visual sensory deprivation during the first months of life strongly affects audio-visual temporal binding later in life. However, given that the TBW narrows during childhood and adolescence in individuals with intact sensory abilities (Hillock-Dunn and Wallace 2012; Hillock et al. 2011; Chen et al. 2016) and that adults undergoing perceptual training demonstrate a narrowing of the TBW (Powers et al. 2009), we conclude that multisensory temporal integration can be influenced by a change in demands also in adults and the lack of differences in TBW between individuals with congenital and acquired anosmia could be attributed to this plasticity. This is further supported by the fact that both congenital and acquired blind individuals exhibit integration patterns linked to a TBW narrowing, namely a low susceptibility to multisensory illusions (Hötting and Röder 2004; Champoux et al. 2011; Occelli et al. 2012). These findings might, however, not generalize to other types of sensory loss, as deaf individuals show an increased susceptibility to multisensory illusions (Heimler et al. 2017; Karns et al. 2012). Moreover, the sensory combinations used to investigate MSI abilities differ among the studied populations: in blind individuals, integration of tactile and auditory stimuli has been assessed, in deaf, integration of tactile and visual stimuli, and we investigated the integration of auditory and visual stimuli. Therefore, direct comparisons are difficult and future studies should attempt to assess MSI abilities of a more diverse set of sensory combinations.

When presented with dynamic, degraded stimuli in the multisensory object identification task, both the anosmia group and the control group displayed multisensory enhancement effects in line with previous results from healthy individuals (Regenbogen et al. 2016; Regenbogen et al. 2018). Specifically, individuals with congenital anosmia demonstrated greater multisensory enhancement effects than healthy controls in the ‘experimental’ setting, i.e., when comparing bimodal to unimodal stimuli; an improvement not demonstrated by individuals with acquired anosmia. In the ‘realistic’ setting, in which the uni- and bimodal conditions were equalized in sensory load by combining the unimodal stimuli with noise in the respective other modality, no significant group differences were demonstrated, albeit individuals with congenital, but not acquired, anosmia demonstrated a statistical trend of improved multisensory enhancement compared to controls. Whether this statistical trend indicates improved multisensory enhancement in individuals with congenital anosmia also in the ‘realistic’ setting cannot be determined based on this study alone and should be further investigated in the future.

The outcome of the two experimental tasks presented slightly heterogeneous results. On one hand, clear group differences were demonstrated in the simultaneity judgement task. On the other hand, only individuals with congenital anosmia demonstrated altered abilities in the object identification task. The difference in results between the two tasks are not uncommon in the sensory loss literature, where studies using multiple experimental tasks can reveal differences between groups of different sensory loss etiology in certain tasks, but no differences in other tasks (Voss et al. 2004). Even though both experimental tasks in the present study investigate audio-visual MSI abilities, they differ in a number of important aspects. First, the experimental tasks focus on different properties of MSI. While the simultaneity judgement assesses multisensory temporal congruency, the object identification is based on the principle of inverse effectiveness. Second, the tasks include stimuli of different complexity: perceptually simple and clear stimuli in the simultaneity judgement, and degraded, dynamic stimuli with semantic information in the object identification task. Lastly, there are also differences in the tasks. The simultaneity judgement is based on the detection of temporal (a)synchronies, knowing exactly which information to search for, while the object identification is based on finding, integrating, and interpreting semantic information which can be present in either (or both) sensory modality. Based on these differences, it is reasonable to argue that separate mechanisms requiring different cognitive processes are involved in the two experimental tasks. Consequently, an enhancement in the perception of audio-visual temporal asynchrony might only be vaguely related to the performance in a task involving semantic comprehension and integration of more complex stimuli. Indeed, independence between the performance on different MSI tasks was recently demonstrated (Odegaard and Shams 2016). Taken together, this could contribute to the understanding of the performance differences between individuals with congenital and acquired anosmia. However, the neural underpinnings of why certain compensatory abilities seem to require a lifelong sensory loss, whereas other alterations emerge also for a sensory loss acquired later in life, need to be determined. A better understanding of the potential cortical reorganization related to olfactory sensory loss is needed to explain the differences in MSI abilities, both within the anosmia group and between individuals with anosmia and controls.

The processes behind the demonstrated multisensory alterations in individuals with anosmia are not yet known. The complete lack of input from one sensory modality, as a result of sensory deprivation, has been shown to extensively alter the neural compositions in multisensory areas normally integrating information from the lost sense with information from other senses. The multisensory areas become more densely populated with neurons processing an intact modality while decreasing the neurons responding to the deprived sense: Monkeys congenitally deprived of visual input display an increased proportion of neurons responding to tactile stimuli in a multisensory cortical area, and a decreased proportion of neurons responding to visual as well as visuo-tactile stimuli, after regaining visual input (Hyvärinen et al. 1981). Similarly, neuroimaging studies in humans have indicated that deaf and blind individuals show increased recruitment of multisensory areas, such as the posterior parietal cortex, when processing stimuli from intact senses (Bavelier et al. 2001). Whether this sensory loss-related recruitment of multisensory cortical areas actually leads to an enhanced integration of the remaining senses is still unclear. Studies on neural populations in multisensory areas in dark-reared cats indicate that the number of neurons responding to audio-tactile stimuli, i.e., the integration of two intact senses, are similar or slightly increased in subcortical (Wallace et al. 2004) as well as cortical multisensory areas (Carriere et al. 2007). Combined, these results suggest that anosmia generates a reorganization of multisensory areas normally processing olfaction, leading to an increased proportion of neurons responding to input from remaining senses and potentially also an increase in multisensory neurons integrating input from remaining senses. Such an olfactory loss-related reorganization of cortical areas governing integration of multisensory stimuli could explain the narrow TBW, here demonstrated by individuals with anosmia, and the improved multisensory enhancement effect of more complex stimuli in individuals with congenital anosmia. Audio-visual temporal integration has been linked to posterior parietal cortex (Zmigrod and Zmigrod 2015) and multisensory enhancement effects of degraded dynamic stimuli, nearly identical to the stimuli in the object identification task, have been demonstrated in the intraparietal sulcus (Regenbogen et al. 2018), a known integration hub of auditory, visual, and tactile stimuli, and an area that also has been linked to integration of olfactory stimuli (Gottfried and Dolan 2003; Boyle et al. 2007; Regenbogen et al. 2017). One could hypothesize that a lack of olfactory input to these parietal, multisensory areas changes the neuronal constellation and promotes a more efficient integration of remaining sensory modalities.

No group differences in response time or number of unanswered trials were demonstrated for either of the two experimental tasks, indicating equal levels of attention for individuals with anosmia and healthy controls. Furthermore, the lack of group differences in the drift diffusion model parameters threshold separation and non-decision time indicate that no differences in response style or non-decision processes (e.g. motor execution), respectively, existed between groups. Therefore, the demonstrated group difference in the two audio-visual integration experiments are unlikely to be attributed to differences in attention or motivation.

In this study, individuals with anosmia demonstrated enhanced audio-visual integration performance. It can be argued that anosmia might also lead to altered auditory or visual unisensory abilities; aspects that were not investigated here and we therefore make no claims in either direction. We do, however, argue that the multisensory differences demonstrated by individuals with anosmia cannot be solely attributed to potential differences in unisensory abilities. In the simultaneity judgment task, the temporal binding of auditory and visual stimuli could potentially be influenced by alterations in unisensory temporal accuracy. That said, temporal precision within the auditory and visual sensory modalities can be as low as 2-3 ms (Zanker and Harris 2002; Gescheider 1966) whereas the demonstrated group difference in TBW was established to be 51 ms. This means that even if potential group differences in unisensory temporal precision exist, they cannot alone explain the demonstrated group difference in multisensory temporal integration. Further, in the thresholding of the object identification task, individuals with anosmia demonstrated a decreased ability to identify unimodal degraded auditory objects compared to healthy controls, as demonstrated by a higher SNR in the individually degraded auditory stimuli. This difference in ability was compensated for in the subsequent multisensory experimental task by using the individually degraded stimuli and thereby equating the individual difficulty level rather than the physical noise level. By assuring that the perceived identification difficulty of all unimodal stimuli was equated across participants, we minimized the influence of potential unisensory group differences and maximized the potential multisensory enhancement according to the principle of inverse effectiveness (Stein and Meredith 1993). Although not statistically significant, it is worth noting that individuals with anosmia demonstrated a statistical trend of enhanced auditory object identification in the “experimental” setting of the multisensory experimental task. As the stimuli were individually tailored, it is unlikely that this is an indication of unisensory differences but could potentially suggest other differences, e.g., higher attention towards auditory stimuli, as the task requires switching of attention towards the sensory modality in which the information is presented. Because individuals with auditory and visual sensory loss often display enhanced performance in remaining sensory modalities, tasks with fixed difficulty levels of stimuli would likely be perceived as easier for these individuals, which in turn would result in lower multisensory enhancements based on the principle of inverse effectiveness. This might partly explain the lower MSI effects demonstrated by deaf and blind individuals in visuo-tactile and audio-tactile tasks, respectively, as compared to non-sensory deprived healthy controls (Hauthal et al. 2014; Collignon, Charbonneau, et al. 2009). However, using complex stimuli with set levels of difficulty would mimic a more ecologically valid scenario in which all individuals are subjected to the same stimuli, rather than stimuli that optimize multisensory enhancement. It would therefore be of relevance in future studies to also assess potential differences in MSI between individuals with anosmia and healthy controls using stimuli with set levels of difficulty.

Lower proportions of simultaneity judgements were overall demonstrated by individuals with anosmia, as compared to healthy controls, in the simultaneity judgement task; albeit both groups’ responses are well within the range of simultaneity judgements from previous reports on healthy individuals (Powers et al. 2009; Stevenson et al. 2012; Scarpina et al. 2016). Independent of whether this difference is spurious or an indication of a potential response bias, the difference is compensated for in the main analysis by not computing the TBW based on a fixed simultaneity judgement but instead, for each individual, using 75 % of peak simultaneity judgement.

In summary, results from two different audio-visual integration tasks indicate an increased sensitivity of multisensory temporal asynchronies in individuals with anosmia, independent of sensory loss etiology, and increased multisensory enhancement in individuals with congenital anosmia. These results suggest the existence of cross-modal compensatory mechanisms in the form of multisensory integration within individuals with olfactory sensory loss; effects more pronounced in those with a lifelong loss. While compensatory mechanisms have long been known to affect processing of isolated senses in individuals with visual and auditory sensory loss, the present study supports the notion that sensory loss enhances the *integration* of remaining senses.

## CONFLICT OF INTEREST

The authors have no conflict of interest to report.

### ACKNOWLEDGEMENT

This work was supported by grants awarded to JNL from the Knut and Alice Wallenberg Foundation (KAW 2012.0141) and the Swedish Research Council (2014-1346).

## DATA, CODE AND STIMULUS AVAILABILITY

Public archiving of data from this study is not permitted by the conditions of our ethics approval. Access to the anonymized data will not be restricted for any individual that seek access to the data with an explicit research purpose. Access to data is granted (without collaboration or legal demands) by the corresponding authors who can be reached by the listed email contacts. Analysis code and experiment/stimulus presentation code are freely and publicly available at https://osf.io/ybfpa/ (DOI 10.17605/OSF.IO/YBFPA). However, copyright restrictions do not permit us to share the full set of stimuli used in this study (specifically the video/audio clips obtained from Shutterstock). Readers may contact the corresponding authors, who are able to provide guidance to stimulus sources. We report how we determined our sample size, all data exclusions, all inclusion/exclusion criteria, whether inclusion/exclusion criteria were established prior to data analysis, all manipulations, and all measures in the study. No part of this study was pre-registered.

## REFERENCES

Alary, F., Duquette, M., Goldstein, R., Elaine Chapman, C., Voss, P., La Buissonnière-Ariza, V. and Lepore, F. 2009. Tactile acuity in the blind: a closer look reveals superiority over the sighted in some but not all cutaneous tasks. Neuropsychologia 47(10), pp. 2037–2043.

Bavelier, D., Brozinsky, C., Tomann, A., Mitchell, T., Neville, H. and Liu, G. 2001. Impact of early deafness and early exposure to sign language on the cerebral organization for motion processing. The Journal of Neuroscience 21(22), pp. 8931–8942.

Bavelier, D., Dye, M.W. and Hauser, P.C. 2006. Do deaf individuals see better? Trends in Cognitive Sciences 10(11), pp. 512–518.

Bavelier, D. and Neville, H.J. 2002. Cross-modal plasticity: where and how? Nature Reviews. Neuroscience 3(6), pp. 443–452.

Bolognini, N., Cecchetto, C., Geraci, C., Maravita, A., Pascual-Leone, A. and Papagno, C. 2012. Hearing shapes our perception of time: temporal discrimination of tactile stimuli in deaf people. Journal of Cognitive Neuroscience 24(2), pp. 276–286.

Boyle, J.A., Frasnelli, J., Gerber, J., Heinke, M. and Hummel, T. 2007. Cross-modal integration of intranasal stimuli: a functional magnetic resonance imaging study. Neuroscience 149(1), pp. 223–231.

Brämerson, A., Johansson, L., Ek, L., Nordin, S. and Bende, M. 2004. Prevalence of olfactory dysfunction: the skövde population-based study. The Laryngoscope 114(4), pp. 733–737.

Carriere, B.N., Royal, D.W., Perrault, T.J., Morrison, S.P., Vaughan, J.W., Stein, B.E. and Wallace, M.T. 2007. Visual deprivation alters the development of cortical multisensory integration. Journal of Neurophysiology 98(5), pp. 2858–2867.

Champoux, F., Collignon, O., Bacon, B.A., Lepore, F., Zatorre, R.J. and Théoret, H. 2011. Early- and late-onset blindness both curb audiotactile integration on the parchment-skin illusion. Psychological Science 22(1), pp. 19–25.

Chen, Y.-C., Shore, D.I., Lewis, T.L. and Maurer, D. 2016. The development of the perception of audiovisual simultaneity. Journal of Experimental Child Psychology 146, pp. 17–33.

Collignon, O., Charbonneau, G., Lassonde, M. and Lepore, F. 2009. Early visual deprivation alters multisensory processing in peripersonal space. Neuropsychologia 47(14), pp. 3236–3243.

Collignon, O., Voss, P., Lassonde, M. and Lepore, F. 2009. Cross-modal plasticity for the spatial processing of sounds in visually deprived subjects. Experimental Brain Research 192(3), pp. 343–358.

Cornell Kärnekull, S., Arshamian, A., Nilsson, M.E. and Larsson, M. 2016. From perception to metacognition: auditory and olfactory functions in early blind, late blind, and sighted individuals. Frontiers in psychology 7, p. 1450.

Cuevas, I., Plaza, P., Rombaux, P., De Volder, A.G. and Renier, L. 2009. Odour discrimination and identification are improved in early blindness. Neuropsychologia 47(14), pp. 3079–3083.

Czarnecki, L., Moberly, A., Fast, C., Turkel, D. and McGann, J. 2018. Multisensory expectations shape olfactory input to the brain. BioRxiv.

Dye, M.W.G., Hauser, P.C. and Bavelier, D. 2009. Is visual selective attention in deaf individuals enhanced or deficient? The case of the useful field of view. Plos One 4(5), p. e5640.

Van Eijk, R.L.J., Kohlrausch, A., Juola, J.F. and van de Par, S. 2008. Audiovisual synchrony and temporal order judgments: effects of experimental method and stimulus type. Perception & Psychophysics 70(6), pp. 955–968.

Frasnelli, J., Collignon, O., Voss, P. and Lepore, F. 2011. Crossmodal plasticity in sensory loss. Progress in Brain Research 191, pp. 233–249.

Frasnelli, J., Schuster, B. and Hummel, T. 2010. Olfactory dysfunction affects thresholds to trigeminal chemosensory sensations. Neuroscience Letters 468(3), pp. 259–263.

Gagnon, L., Vestergaard, M., Madsen, K., Karstensen, H.G., Siebner, H., Tommerup, N., Kupers, R. and Ptito, M. 2014. Neural correlates of taste perception in congenital olfactory impairment. Neuropsychologia 62, pp. 297–305.

Gelman, A. and Rubin, D.B. 1992. Inference from Iterative Simulation Using Multiple Sequences. Statistical Science 7(4), pp. 457–472.

Gescheider, G.A. 1966. Resolving of successive clicks by the ears and skin. Journal of experimental psychology 71(3), pp. 378–381.

Goldreich, D. and Kanics, I.M. 2003. Tactile acuity is enhanced in blindness. The Journal of Neuroscience 23(8), pp. 3439–3445.

Gottfried, J.A. and Dolan, R.J. 2003. The nose smells what the eye sees: crossmodal visual facilitation of human olfactory perception. Neuron 39(2), pp. 375–386.

Gougoux, F., Lepore, F., Lassonde, M., Voss, P., Zatorre, R.J. and Belin, P. 2004. Neuropsychology: pitch discrimination in the early blind. Nature 430(6997), p. 309.

Hauthal, N., Debener, S., Rach, S., Sandmann, P. and Thorne, J.D. 2014. Visuo-tactile interactions in the congenitally deaf: a behavioral and event-related potential study. Frontiers in Integrative Neuroscience 8, p. 98.

Heimler, B., Baruffaldi, F., Bonmassar, C., Venturini, M. and Pavani, F. 2017. Multisensory interference in early deaf adults. Journal of deaf studies and deaf education 22(4), pp. 422–433.

Hillock, A.R., Powers, A.R. and Wallace, M.T. 2011. Binding of sights and sounds: age-related changes in multisensory temporal processing. Neuropsychologia 49(3), pp. 461–467.

Hillock-Dunn, A. and Wallace, M.T. 2012. Developmental changes in the multisensory temporal binding window persist into adolescence. Developmental Science 15(5), pp. 688–696.

Hummel, T., Futschik, T., Frasnelli, J. and Hüttenbrink, K.-B. 2003. Effects of olfactory function, age, and gender on trigeminally mediated sensations: a study based on the lateralization of chemosensory stimuli. Toxicology Letters 140-141, pp. 273–280.

Hummel, T., Kobal, G., Gudziol, H. and Mackay-Sim, A. 2007. Normative data for the “Sniffin’ Sticks” including tests of odor identification, odor discrimination, and olfactory thresholds: an upgrade based on a group of more than 3,000 subjects. European Archives of Oto-Rhino-Laryngology 264(3), pp. 237–243.

Hummel, T., Whitcroft, K.L., Andrews, P., Altundag, A., Cinghi, C., Costanzo, R.M., Damm, M., Frasnelli, J., Gudziol, H., Gupta, N., Haehner, A., Holbrook, E., Hong, S.C., Hornung, D., Hüttenbrink, K.B., Kamel, R., Kobayashi, M., Konstantinidis, I., Landis, B.N., Leopold, D.A., Macchi, A., Miwa, T., Moesges, R., Mullol, J., Mueller, C.A., Ottaviano, G., Passali, G.C., Philpott, C., Pinto, J.M., Ramakrishnan, V.J., Rombaux, P., Roth, Y., Schlosser, R.A., Shu, B., Soler, G., Stjärne, P., Stuck, B.A., Vodicka, J. and Welge-Luessen, A. 2017. Position paper on olfactory dysfunction. Rhinology.

Hyvärinen, J., Hyvärinen, L. and Linnankoski, I. 1981. Modification of parietal association cortex and functional blindness after binocular deprivation in young monkeys. Experimental Brain Research 42(1), pp. 1–8.

Hötting, K. and Röder, B. 2004. Hearing cheats touch, but less in congenitally blind than in sighted individuals. Psychological Science 15(1), pp. 60–64.

Karns, C.M., Dow, M.W. and Neville, H.J. 2012. Altered cross-modal processing in the primary auditory cortex of congenitally deaf adults: a visual-somatosensory fMRI study with a double-flash illusion. The Journal of Neuroscience 32(28), pp. 9626–9638.

Landis, B.N., Konnerth, C.G. and Hummel, T. 2004. A study on the frequency of olfactory dysfunction. The Laryngoscope 114(10), pp. 1764–1769.

Landis, B.N., Scheibe, M., Weber, C., Berger, R., Brämerson, A., Bende, M., Nordin, S. and Hummel, T. 2010. Chemosensory interaction: acquired olfactory impairment is associated with decreased taste function. Journal of Neurology 257(8), pp. 1303–1308.

Legge, G.E., Madison, C., Vaughn, B.N., Cheong, A.M.Y. and Miller, J.C. 2008. Retention of high tactile acuity throughout the life span in blindness. Perception & Psychophysics 70(8), pp. 1471–1488.

Lessard, N., Paré, M., Lepore, F. and Lassonde, M. 1998. Early-blind human subjects localize sound sources better than sighted subjects. Nature 395(6699), pp. 278–280.

Lewald, J. 2002. Vertical sound localization in blind humans. Neuropsychologia 40(12), pp. 1868–1872.

Levänen, S. and Hamdorf, D. 2001. Feeling vibrations: enhanced tactile sensitivity in congenitally deaf humans. Neuroscience Letters 301(1), pp. 75–77.

Lundström, J.N., Boesveldt, S. and Albrecht, J. 2011. Central processing of the chemical senses: an overview. ACS Chemical Neuroscience 2(1), pp. 5–16.

Lundström, J.N., Regenbogen, C., Ohla, K. and Seubert, J. 2018. Prefrontal control over occipital responses to crossmodal overlap varies across the congruency spectrum. Cerebral Cortex.

Merabet, L.B. and Pascual-Leone, A. 2010. Neural reorganization following sensory loss: the opportunity of change. Nature Reviews. Neuroscience 11(1), pp. 44–52.

Meredith, M.A., Nemitz, J.W. and Stein, B.E. 1987. Determinants of multisensory integration in superior colliculus neurons. I. Temporal factors. The Journal of Neuroscience 7(10), pp. 3215–3229.

Murray, M.M., Lewkowicz, D.J., Amedi, A. and Wallace, M.T. 2016. Multisensory Processes: A Balancing Act across the Lifespan. Trends in Neurosciences 39(8), pp. 567–579.

Navarro, D.J. and Fuss, I.G. 2009. Fast and accurate calculations for first-passage times in Wiener diffusion models. Journal of mathematical psychology 53(4), pp. 222–230.

Occelli, V., Bruns, P., Zampini, M. and Röder, B. 2012. Audiotactile integration is reduced in congenital blindness in a spatial ventriloquism task. Neuropsychologia 50(1), pp. 36–43.

Odegaard, B. and Shams, L. 2016. The brain’s tendency to bind audiovisual signals is stable but not general. Psychological Science 27(4), pp. 583–591.

Perez, F. and Granger, B.E. 2007. IPython: A System for Interactive Scientific Computing. Computing in science & engineering 9(3), pp. 21–29.

Pirozzo, S., Papinczak, T. and Glasziou, P. 2003. Whispered voice test for screening for hearing impairment in adults and children: systematic review. BMJ (Clinical Research Ed.) 327(7421), p. 967.

Porada, D.K., Regenbogen, C., Seubert, J., Freiherr, J. and Lundström, J.N. 2019. Multisensory enahncement of odor object processing in primary olfactory cortex.

Powers, A.R., Hillock, A.R. and Wallace, M.T. 2009. Perceptual training narrows the temporal window of multisensory binding. The Journal of Neuroscience 29(39), pp. 12265–12274.

Putzar, L., Goerendt, I., Lange, K., Rösler, F. and Röder, B. 2007. Early visual deprivation impairs multisensory interactions in humans. Nature Neuroscience 10(10), pp. 1243–1245.

Ratcliff, R. 1978. A theory of memory retrieval. Psychological Review 85(2), pp. 59–108.

Regenbogen, C., Axelsson, J., Lasselin, J., Porada, D.K., Sundelin, T., Peter, M.G., Lekander, M., Lundström, J.N. and Olsson, M.J. 2017. Behavioral and neural correlates to multisensory detection of sick humans. Proceedings of the National Academy of Sciences of the United States of America 114(24), pp. 6400–6405.

Regenbogen, C., Johansson, E., Andersson, P., Olsson, M.J. and Lundström, J.N. 2016. Bayesian-based integration of multisensory naturalistic perithreshold stimuli. Neuropsychologia 88, pp. 123–130.

Regenbogen, C., Seubert, J., Johansson, E., Finkelmeyer, A., Andersson, P. and Lundström, J.N. 2018. The intraparietal sulcus governs multisensory integration of audiovisual information based on task difficulty. Human Brain Mapping 39(3), pp. 1313–1326.

Reichert, J.L. and Schöpf, V. 2018. Olfactory loss and regain: lessons for neuroplasticity. The Neuroscientist 24(1), pp. 22–35.

Rombaux, P., Huart, C., De Volder, A.G., Cuevas, I., Renier, L., Duprez, T. and Grandin, C. 2010. Increased olfactory bulb volume and olfactory function in early blind subjects. Neuroreport 21(17), pp. 1069–1073.

Röder, B., Teder-Sälejärvi, W., Sterr, A., Rösler, F., Hillyard, S.A. and Neville, H.J. 1999. Improved auditory spatial tuning in blind humans. Nature 400(6740), pp. 162–166.

Scarpina, F., Migliorati, D., Marzullo, P., Mauro, A., Scacchi, M. and Costantini, M. 2016. Altered multisensory temporal integration in obesity. Scientific reports 6, p. 28382.

Singh, A.K., Phillips, F., Merabet, L.B. and Sinha, P. 2018. Why Does the Cortex Reorganize after Sensory Loss? Trends in Cognitive Sciences 22(7), pp. 569–582.

Small, D.M. 2012. Flavor is in the brain. Physiology & Behavior 107(4), pp. 540–552.

Snellen, H. 1862. Probebuchstaben zur Bestimmung der Sehschärfe. Utrecht.

Sorokowska, A., Sorokowski, P., Karwowski, M., Larsson, M. and Hummel, T. 2018. Olfactory perception and blindness: a systematic review and meta-analysis. Psychological research.

Statistiska centralbyrån, S.C.B. 2018. The labour market situation for people with disabilities 2017. Örebro, Sweden: Statistiska centralbyrån.

Stein, B.E. and Meredith, M.A. 1993. The merging of the senses. Cambridge, MA, US: MIT Press.

Stein, B.E. and Stanford, T.R. 2008. Multisensory integration: current issues from the perspective of the single neuron. Nature Reviews. Neuroscience 9(4), pp. 255–266.

Stevenson, R.A., Ghose, D., Fister, J.K., Sarko, D.K., Altieri, N.A., Nidiffer, A.R., Kurela, L.R., Siemann, J.K., James, T.W. and Wallace, M.T. 2014. Identifying and quantifying multisensory integration: a tutorial review. Brain Topography 27(6), pp. 707–730.

Stevenson, R.A., Siemann, J.K., Schneider, B.C., Eberly, H.E., Woynaroski, T.G., Camarata, S.M. and Wallace, M.T. 2014. Multisensory temporal integration in autism spectrum disorders. The Journal of Neuroscience 34(3), pp. 691–697.

Stevenson, R.A., Zemtsov, R.K. and Wallace, M.T. 2012. Individual differences in the multisensory temporal binding window predict susceptibility to audiovisual illusions. Journal of Experimental Psychology. Human Perception and Performance 38(6), pp. 1517–1529.

Wallace, M.T., Perrault, T.J., Hairston, W.D. and Stein, B.E. 2004. Visual experience is necessary for the development of multisensory integration. The Journal of Neuroscience 24(43), pp. 9580–9584.

Wallace, M.T. and Stevenson, R.A. 2014. The construct of the multisensory temporal binding window and its dysregulation in developmental disabilities. Neuropsychologia 64, pp. 105–123.

Wesson, D.W. and Wilson, D.A. 2010. Smelling sounds: olfactory-auditory sensory convergence in the olfactory tubercle. The Journal of Neuroscience 30(8), pp. 3013–3021.

Wiecki, T.V., Sofer, I. and Frank, M.J. 2013. HDDM: Hierarchical Bayesian estimation of the Drift-Diffusion Model in Python. Frontiers in Neuroinformatics 7, p. 14.

Voss, A., Nagler, M. and Lerche, V. 2013. Diffusion models in experimental psychology: a practical introduction. Experimental psychology 60(6), pp. 385–402.

Voss, P. 2013. Sensitive and critical periods in visual sensory deprivation. Frontiers in psychology 4, p. 664.

Voss, P., Lassonde, M., Gougoux, F., Fortin, M., Guillemot, J.-P. and Lepore, F. 2004. Early- and late-onset blind individuals show supra-normal auditory abilities in far-space. Current Biology 14(19), pp. 1734–1738.

Voss, P. and Zatorre, R.J. 2012. Occipital cortical thickness predicts performance on pitch and musical tasks in blind individuals. Cerebral Cortex 22(11), pp. 2455–2465.

Vroomen, J. and Keetels, M. 2010. Perception of intersensory synchrony: a tutorial review. Attention, perception & psychophysics 72(4), pp. 871–884.

Zanker, J.M. and Harris, J.P. 2002. On temporal hyperacuity in the human visual system. Vision Research 42(22), pp. 2499–2508.

Zmigrod, L. and Zmigrod, S. 2016. On the temporal precision of thought: individual differences in the multisensory temporal binding window predict performance on verbal and nonverbal problem solving tasks. Multisensory research 29(8), pp. 679–701.

Zmigrod, S. and Zmigrod, L. 2015. Zapping the gap: Reducing the multisensory temporal binding window by means of transcranial direct current stimulation (tDCS). Consciousness and Cognition 35, pp. 143–149.

Zwiers, M.P., Van Opstal, A.J. and Cruysberg, J.R. 2001. A spatial hearing deficit in early-blind humans. The Journal of Neuroscience 21(9), p. RC142: 1–5.

